# *LYS3* encodes a prolamin-box-binding transcription factor that controls embryo growth in barley and wheat

**DOI:** 10.1101/2019.12.18.880955

**Authors:** Beata Orman-Ligeza, Philippa Borrill, Tansy Chia, Marcella Chirico, Jaroslav Doležel, Sinead Drea, Miroslava Karafiátová, Nicole Schatlowski, Charles U. Solomon, Burkhard Steuernagel, Brande B. H. Wulff, Cristobal Uauy, Kay Trafford

**Author notes:** Corresponding author. Kay Trafford, The National Institute of Agricultural Botany, Huntingdon Road, Cambridge, Cambridgeshire CB3 0LE, UK. Present addresses: Nicole Schatlowski: Wellcome Genome Campus, Hinxton, CB10 1SA, UK. Abbreviations:* Days after flowering (DAF), DNA binding with one zinc finger (DOF), prolamin-box binding factor (PBF), targeting induced local lesions in genomes (TILLING), sorting intolerant from tolerant (SIFT).

## Abstract

Mutations at the *LYS3* locus in barley have multiple effects on grain development, including an increase in embryo size and a decrease in endosperm starch content. The gene underlying *LYS3* was identified by genetic mapping and mutations in this gene were identified in all four barley *lys3* alleles. *LYS3* encodes a transcription factor called Prolamin Binding Factor (PBF). Its role in controlling embryo size was confirmed using wheat TILLING mutants. To understand how *PBF* controls embryo development, we studied its spatial and temporal patterns of expression in developing grains. The *PBF* gene is expressed in both the endosperm and the embryos, but the timing of expression in these organs differs. *PBF* expression in wild-type embryos precedes the onset of embryo enlargement in *lys3* mutants, suggesting that *PBF* suppresses embryo growth. We predicted the down-stream target genes of *PBF* in wheat and found them to be involved in a wide range of biological processes, including organ development and starch metabolism. Our work suggests that *PBF* may influence embryo size and endosperm starch synthesis via separate gene control networks.

**HIGHLIGHTS:**

- *LYS3* encodes a transcription factor called Prolamin Binding Factor (PBF) that is expressed in grains only.
- Wheat and barley *LYS3*/*PBF* mutants have enlarged embryos suggesting that this gene suppresses embryo growth.
- The down-stream target genes of *PBF* in wheat are predicted to be involved in a wide range of biological processes including organ development and starch metabolism.

## A. Introduction

In the 1960-70s, in an attempt to improve the lysine content of barley (*Hordeum vulgare* L.) for animal feed, mutagenized barley germplasm at the Risø National Laboratory, Denmark, was screened for lysine content and a number of high-lysine (*lys*) mutants were identified (Doll *et al*., 1974; Doll, 1976). The *lys* mutant with the highest lysine content (44 percent higher than wild type) was Risø1508 (*lys3a*) (Ingversen *et al*., 1973; Mossberg, 1969). In addition to *lys3a*, there are three other barley lines with mutations at the same locus: Risø18 (*lys3b*), Risø19 (*lys3c*) and M1460 (*lys3d*) (Aastrup, 1983; Munck, 1992) (Supplementary Table S1). All four *lys3* mutations are recessive.

Studies have shown that the endosperm of *lys3* mutants, like that of most other high-lysine barley mutants, contains less starch and lysine-poor protein (hordein), but has more lysine-rich protein and free lysine (Brandt, 1976; Ingversen *et al*., 1973). However, uniquely amongst high-lysine barley mutants, the embryos of all four *lys3* mutants are larger than normal (Tallberg, 1977, Deggerdal *et al*., 1986, Cook *et al*., 2018). Most of the barley *lys* mutants, including those known to have lesions in genes encoding components of the starch biosynthesis pathway (Trafford and Fincher, 2014), have reduced embryo weight (as well as shrivelled-endosperm and reduced starch content; Cook *et al*., 2018). The enlargement of the embryos in *lys3* mutants is therefore a specific response to mutation at the *LYS3* locus and not due to the diversion of resources from the shrivelled endosperm to the embryo. As well as increased size, the embryos of *lys3* mutants have higher-than-normal starch content (Deggerdal *et al*., 1986; Olsen *et al*., 1984) and larger-than-normal cells in the scutellum (Olsen *et al*., 1984; Deggerdal *et al*., 1986). The mutant embryos also show reduced dormancy (Cook *et al*., 2018).

Risø1508 (*lys3a*) has been the subject of a breeding programme to improve the nutritional quality of barley, particularly for pig feed (Munck and Jespersen, 2009). Animal feeding trials, using the Risø1508-derived lines Piggy and Lysimax, showed that *lys3a* mutant lines are a more effective source of protein for maximum growth rate in pigs than the wild-type barley varieties from which they are derived (Munck, 1972; Mortensen *et al*., 1988; Gabert *et al*., 1995; Gabert *et al*., 1996). However, the success of Risø1508-derived barley lines as animal feed is hampered by their reduced starch content which leads to low grain weight and consequently to low yield.

To separate the favourable (nutritional enhancements) and unfavourable (yield depression) traits associated with *lys3* mutations, it is necessary to identify the gene responsible and then to understand how it functions in developing embryos and endosperm. We report here the identification of the *LYS3* gene by genetic mapping using embryo size as the selection phenotype. Whilst our work on this was in progress, the gene underlying the *LYS3* locus was independently identified by Moehs *et al*. (2019) using variation in hordein content as the selection phenotype. Both studies agree that *LYS3* encodes a previously-identified and well-studied transcription factor in barley: <u>p</u>rolamin-box <u>b</u>inding factor (PBF). To understand PBF function, we studied its gene expression patterns in barley and found that contrary to previous reports (Mena *et al*., 1998; Mena *et al*., 2002), *PBF* is expressed in developing embryos as well as in the endosperm. The emphasis of our current work is to identify the down-stream target genes of PBF and here we present our initial studies of its predicted targets in wheat.

## 2. Materials and methods

### 2.1. Barley germplasm

Grains of Bomi, Morex and Risø1508 were obtained from the Germplasm Resources Unit, John Innes Centre, Norwich, UK and Risø18, Risø19, M1460 and Minerva were kindly supplied by Birthe Møller Jespersen, University of Copenhagen, Denmark.

### 2.2. Plant growth

For mapping experiments, individual grains were germinated in Petri dishes on moist filter paper. After over-night incubation at 4 °C, plates were transferred to room temperature. When roots and shoots were established, each seedling was transplanted into a 1 L pot containing Levington M2 compost (Scotts Professional, Ipswich, UK) and grown in a glasshouse. In winter, additional lighting was provided by sodium lamps for 16 h per day and temperatures were maintained between 15 °C (night) and 20 °C (day). In summer, plants were grown in a glasshouse under ambient conditions.

Wheat TILLING mutants were sown directly into M2 compost, incubated at 4 °C for 3 days and then transferred to a glasshouse with a 22-hour photoperiod and temperatures of 21 °C (night) and 18 °C (day). Supplementary lighting was provided by a mixture of high-pressure sodium lamps and both far red and white LED lights (Conviron, Winnipeg, US).

### 2.3. Analysis of grain and embryo development

Anthesis occurred whilst the ear was enveloped in the flag leaf so the exact day of anthesis was difficult to determine without damaging the developing spike. Accordingly, flowering time was defined as the day on which the awns of the developing ear protruded more than 1 cm above the leaf sheath and grain/embryo age was measured in days after flowering (DAF).

### 2.4. DNA extraction

For barley genotyping, DNA was extracted as follows: leaf material was harvested at seedling stage to 1.5-ml tubes each containing a 5-mm diameter steel ball and frozen at -80 °C. Frozen leaf material was homogenised using a Geno/Grinder (SPEX SamplePrep LLC) and then 600 µl of extraction buffer (200 mM Tris pH 7.5, 250 mM NaCl, 25 mM EDTA, 0.5% [w/v] SDS) was added. The homogenized leaf material was incubated at 65 °C for one hour. Leaf debris was pelleted by centrifugation and the supernatant was transferred to a fresh 1.5-ml tube, mixed with an equal volume of isopropanol and centrifuged. The pellet was washed with 500 ml 70% [v/v] ethanol and resuspended in 200 µl H_2_O. DNA was quantified using a spectrometer (NanoDrop 1000, Thermo Scientific) and the concentration adjusted to 10 ng/µl.

For wheat genotyping, DNA was extracted from seedling leaf material using the method of Fulton *et al*. (1995).

### 2.5. Chromosome sequencing

For each barley line, suspensions of intact mitotic metaphase chromosomes were prepared from synchronized root tip cells of barley seedlings as described by Lysák *et al*. (1999). Chromosomes in suspension were stained with 2 µg.ml^-1^ DAPI (4’, 6-diamidino-2-phenylindole) and chromosome 5H was sorted using a FACSAria II SORP flow cytometer and sorter (Becton Dickinson Immunocytometry Systems, San José, USA). Purity in the sorted 5H fractions was determined microscopically after FISH with a probe for GAA microsatellite (Kubaláková *et al.,* 2003). DNA of the sorted chromosomes was purified and amplified by multiple displacement amplification according to Šimková *et al*. (2008). Three independent amplification products were combined in each cultivar to reduce amplification bias.

The samples of pooled amplified chromosomal DNA were subjected to Illumina HiSeq2500 sequencing (The Genome Analysis Centre, Norwich, UK; now the Earlham Institute). Paired-end read size was 250 bp for the wild-type samples and 125 bp for the mutant samples. The total number of paired-reads obtained was 172,974,201 (Bomi), 252,692,421 (Risø1508) and 226,936,729 (Risø19). Data were submitted to the European Nucleotide Archive (www.ebi.ac.uk) with accession number PRJEB33709.

### 2.6. Genotyping the mapping population

The Bomi chromosome sequencing data was assembled using the published Morex genome sequence (as described below for MutChromSeq). Genes flanking the *LYS3* locus with SNPs between Bomi and Morex were selected for KASP marker design. For primer sequences see Supplementary Table S2A. Genotyping was performed on a Quant Studio 7 (Applied Biosystems) using KASP technology (LGC) following the manufacturers’ instructions.

### 2.7. Determination of embryo size

For fine mapping in barley, the embryo phenotype of recombinant lines was visually determined and categorized as normal or large embryo. For determination of relative embryo dry weight, all barley or wheat plants for each experiment were grown together in one batch and grains were harvested at maturity. To extract the embryos, grains were soaked overnight in sterile water at 4 °C in the dark and then dissected into embryo and non-embryo portions. Both portions were dried to constant weight by heating in an oven at 65 °C for 3 days. Relative embryo weight (%) = (embryo dry weight / (embryo + non-embryo dry weight)) x 100.

### 2.8. Identification of a candidate gene using MutChromSeq

The chromosome sequencing data was analysed using the MutChromSeq pipeline according to Steuernagel *et al*. (2017). The raw data were trimmed using sickle (https://github.com/najoshi/sickle) with default parameters. Trimmed data from wild types Bomi and Minerva were assembled using CLC Assembly Cell (https://www.qiagenbioinformatics.com/products/clc-assembly-cell/), version 5.1 with default parameters. Trimmed data from mutants and wild types were mapped to wild-type assemblies using bwa sampe (Li and Durbin, 2009), version 0.7.12 with default parameters. Further processing of mappings was done using samtools (Li *et al*., 2009) version 0.1.19. For running MutChromSeq, we used release 2 (https://github.com/steuernb/MutChromSeq/releases/tag/2), which introduces the filtering of contigs using a mapping interval, as described in Dracatos *et al*. (2019). These sequences and those of the wild-type controls have been submitted to GenBank (accession numbers: MN715383, MN715384, MN715385, MN715386, MN715387).

### 2.9. Analysis of barley PBF/LYS3 genes

The SNP mutations in the *PBF/LYS3* genes that were identified by chromosome sequencing were confirmed by PCR amplification and Sanger sequencing. The primer sequences and PCR conditions are given in Supplementary Table S2B. Mutations in the mapping lines were confirmed using KASP assays (Supplementary Table S2C).

### 2.10. Large-embryo TILLING mutants of wheat

Lines of the hexaploid wheat cultivar Cadenza were selected from the *in silico* wheat TILLING resource (www.wheat-tilling.com; Krasileva et al., 2017) (Supplementary Table S2D). The supplied seed was sown, pairs of homozygous mutant lines were selected and cross-pollinated, and the resulting F_1_ plants were confirmed to be heterozygous. A homozygous triple mutant was constructed by crossing Cadenza0903 (*TaPBF-B1*) to Cadenza0904 (*TaPBF-D1*) and homozygous F_2_ double mutants were then crossed to Cadenza1807 or Cadenza1553 (*TaPBF-A1* mutants).

### 2.11. Analysis of expression by Reverse Transcription PCR

Tissue was harvested, weighed, frozen in liquid nitrogen and stored at -20 °C. Total RNA was extracted using Tri Reagent (Sigma Aldrich, UK) according to the manufacturer’s instructions, its concentration was measured with a spectrometer (NanoDrop 1000, Thermo Scientific) and it was stored at -80 °C. An aliquot containing 4 μg RNA was treated for 45 min with 2 μl of DNase RQ1 (1 μg/μl) at 37 °C (Promega, UK), and purified using an RNAeasy spin column (Qiagen, UK). cDNA was prepared from 0.5 μg RNA using a SuperScript III reverse transcriptase kit and oligo(dT)18 primers (Thermo Fisher Scientific, UK) according to the manufacturer’s instructions and stored at -20 °C prior to PCR. The primer sequences and PCR conditions are given in Supplementary Table S2E.

### 2.12. Analysis of expression by in situ hybridization

Developing barley grains were harvested and processed for mRNA *in situ* hybridization as described in Drea *et al*., (2005) and Opanowicz *et al.,* (2010). The probe template consisted of a *PBF* cDNA fragment amplified with gene-specific primers from a barley grain cDNA library (primer sequences are given in Supplementary Table S2F). For amplification of the antisense probe, a T7 polymerase promoter sequence was attached to the 3’ end of the reverse primer. For amplification of the sense probe, a T7 promoter sequence was attached to the 5’ end of the forward primer. The amplified products were transcribed *in vitro* with T7 RNA polymerase (Bioline, UK).

### 2.13. Downstream targets of wheat PBF

The promotor regions (−2 kb) of 29 wheat starch biosynthesis genes were searched for sequences conforming to the consensus prolamin and pyrimidine binding sequences (TGTAAAG and CCTTTT, respectively). The predicted TaPBF downstream target genes were investigated using the RefSeqv1.0 wheat genome assembly and associated wheat gene networks. One network (referred to as the GENIE3 network) had been constructed to predict transcription factor targets using 850 RNA-seq samples (Ramírez-González *et al*. (2018) and four separate co-expression (Weight Gene Correlation Network Analysis, WGCNA) networks for grain, leaf, root and spike tissues were used to identify co-expressed genes (Ramírez-González et al., 2018). Gene ontology term enrichment analysis was performed as described in Ramírez-González *et al*. (2018) using GOseq (Young et al., 2010).

## 3. Results

### 3.1. Fine mapping LYS3 identified a 64-gene region on chromosome arm 5HL controlling embryo size

Our previous work suggested that the *LYS3* gene in barley controls embryo size. To test this and to identify the *LYS3* gene, we developed a mapping population by crossing each of the three Risø *lys3* large-embryo mutants to Morex, a cultivar with normal embryo size and with available genome sequence data. Following self-pollination, the F_2_ grains were genotyped with polymorphic markers in genes close to the *LYS3* locus on chromosome arm 5HL (Franckowiak, 1997). Lines with chromosomal recombination points close to *LYS3* were selected and allowed to self-pollinate. No lines with recombination events close to *lys3* were found for progeny from the Risø18 x Morex crosses and so further work involved progeny from crosses between Risø19 and Risø1508 only. Homozygous recombinant lines (and controls) were selected from the progeny for further genotypic and phenotypic analysis.

To design new markers for genotyping within the region of interest, we purified by flow cytometric sorting the 5H chromosomes from both *lys3* mutants and the parental control cultivar, Bomi, and we sequenced DNA amplified from them using Illumina HiSeq2500. The sequencing data was assembled using the MutChromSeq pipeline according to Steuernagel *et al*. (2017), and single nucleotide polymorphisms (SNPs) in the region of interest were identified. These SNPs were used to design KASP primers for further genotypic analysis (Supplementary Table S2).

To phenotype the homozygous recombinant lines, we first assessed embryo size visually (Table 1; Fig. 1A). The phenotype of selected critical lines was also assessed by measurement of relative embryo weight and β-amylase activity (Fig. 1B). In our previous work, we showed that all four *lys3* mutants had increased absolute and relative embryo weights (Cook *et al*., 2018). When tested again in different growth conditions (Fig. 2A), although the relative embryo size was consistently increased for all *lys3* mutants, the increase in absolute embryo weight was significant for Risø1508 and M1460 only. For this reason, we used the relative embryo weight to phenotype the critical mapping lines. We also used β-amylase activity to phenotype these critical lines because it is known to be strongly affected by *lys3* mutations (Allison, 1978).

**Figure 1.**
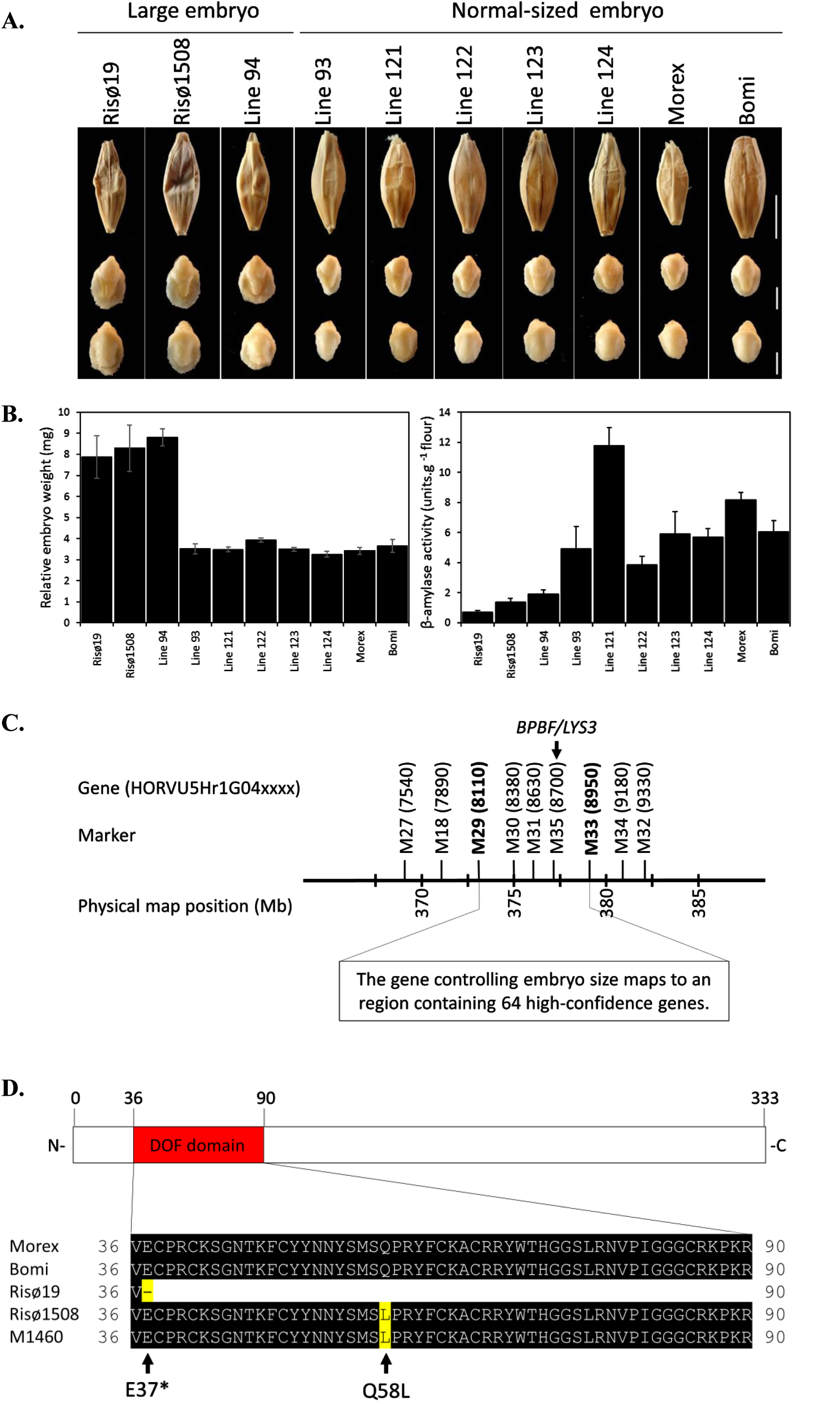
Map-based cloning of the gene controlling embryo size. A. Representative images of grains and embryos from the lines described in Table 1 are shown. Top row = dry, mature grains. Middle row = excised embryos, abaxial surface uppermost. Lower row = excised embryos, adaxial surface uppermost. The scale bars are 5 mm for grains and 2 mm for embryos. B. Relative embryo weight and β-amylase activity were determined for selected lines shown in Table 1. The values for relative embryo weight are the means ± SE for 5 plants and 10 grains were measured per plant. The values for β-amylase activity are the means ± SE for three plants. One grain from the middle of the ear was assayed per plant. All assays were triplicated. C. The gene controlling embryo size maps to a region on chromosome arm 5HL. The two markers/genes flanking this region are shown in bold. The *LYS3*/*PBF* gene controlling embryo size was identified as HORVU5Hr1G048700. Gene names are abbreviated: xxxx = HORVU5Hr1G04xxxx. D. An alignment of PBF amino acid sequences showing the positions of the DOF domain and the SNP mutations. (Risø18 lacks *PBF* and is therefore not shown).

**Figure 2.**
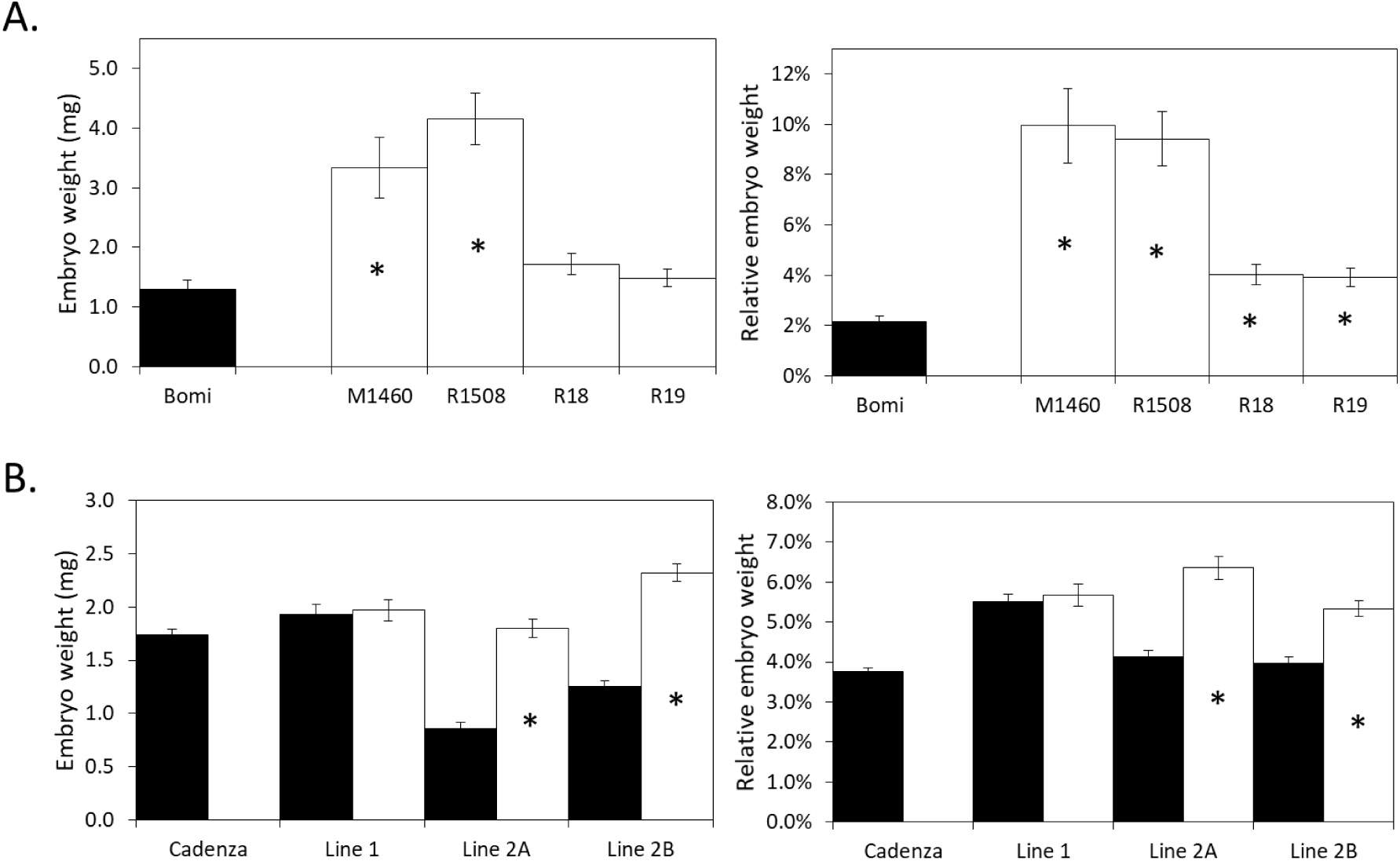
*PBF* affects embryo weight in barley and wheat. The absolute and relative dry weights of the embryo and non-embryo grain parts of wild-types (black bars) and mutants (white bars) are shown. Mean values ±SE are for 11 to 13 (barley) or 25 (wheat) replicate biological samples. Each sample contained 10 grains. Values for mutants that are significantly different from the values for their relevant wild-type controls (Students t-test, p<0.05) are indicated by an asterisk. A. Barley mutants. Risø1508, Risø18 and Risø19 and M1460 are *lys3* mutants. Bomi is the wild-type parent of the three Risø mutants. B. Wheat TILLING mutants. Lines are as described in Figure 3.

**Table 1.** Fine-mapping LYS3 in barley. The genotypes of genes close to LYS3 are shown: A = wild type (Morex), B = Bomi and lys3 mutant. The embryo-size phenotype was assessed visually: WT = wild type (normal-sized embryos), MUT = mutant (large embryos). The two genes defining the *LYS3* region are shown in bold. The *PBF* gene, HORVU5Hr1G048700 is underlined.

| Line | *lys3* parent | Visual embryo phenotype | Number of plants analysed | Genotype (Gene ID = HORVU5Hr1G04xxxx) |  |  |  |  |  |  |  |  |
| --- | --- | --- | --- | --- | --- | --- | --- | --- | --- | --- | --- | --- |
|  |  |  |  | 7540 | 7890 | **8110** | 8380 | 8630 | <u>8700</u> | **8950** | 9180 | 9330 |
| Morex |  | WT (control) |  | A | A | A | A | A | A | A | A | A |
| Line 92 | Risø19 | WT | 2 | B | B | A | A | A | A | A | A | A |
| Line 93 | Risø1508 | WT | 3 | B | B | B | A | A | A | A | A | A |
| Line 123 | Risø1508 | WT | 2 | B | B | B | A | A | A | A | A | A |
| Line 94 | Risø19 | MUT | 1 | B | B | B | B | B | B | A | A | A |
| Line 125 | Risø1508 | MUT | 1 | B | B | B | B | B | B | B | B | A |
| Line 115 | Risø1508 | WT | 1 | A | A | A | A | A | A | A | B | B |
| Line 121 | Risø1508 | WT | 8 | A | A | A | A | A | A | B | B | B |
| Line 122 | Risø19 | WT | 6 | A | A | A | A | A | A | B | B | B |
| Line 124 | Risø1508 | WT | 5 | A | A | A | A | A | A | B | B | B |
| Line 116 | Risø19 | MUT | 10 | A | A | B | B | B | B | B | B | B |
| Line 117 | Risø19 | MUT | 3 | A | A | B | B | B | B | B | B | B |
| Risø19 |  | MUT (control) |  | B | B | B | B | B | B | B | B | B |
| Risø1508 |  | MUT (control) |  | B | B | B | B | B | B | B | B | B |
| Bomi |  | WT |  | B | B | B | B | B | B | B | B | B |

The visual phenotyping together with the genotyping data suggested that the gene that is responsible for the large-embryo phenotype lies between HORVU5Hr1G048110 and HORVU5Hr1G048950 (Table 1, Fig. 1A). Quantitative measurement of relative embryo size in critical lines confirmed the visual phenotyping results (Fig. 1B). The embryo weights for lines visually assessed as ‘mutant’ were significantly greater than the weights of those assessed as ‘wild-type’ (p< 0.05, Student’s *t*-test). The phenotyping results for β-amylase activity were less clear due to a large amount of variation between the values within each group. Although collectively, the large-embryo lines had lower β-amylase activity than the wild-type lines, not all of the comparative differences between individual mutant and wild type lines were statistically significant (Fig. 1B).

### 3.2. LYS3 encodes the transcription factor, PBF

To identify the gene responsible for controlling embryo size within the region of interest identified by mapping, we isolated and sequenced the 5H chromosome of the two other *lys3* mutants, Risø18 and M1460 and from an additional wild-type line, Minerva which is the parent of M1460. This sequencing information was combined with that of the three lines sequenced previously and bioinformatically assessed using a method based on MutChromSeq (a method that was used previously e.g. for the identification of disease-resistance genes in mutant barleys; Sanchez-Martin *et al*., 2016). Only one gene in the region of interest (Fig. 1C) had mutations in all four mutant lines relative to the wild-type controls (Fig. 1D) and that was HORVU5Hr1G048700 which encodes the PBF transcription factor.

The *PBF* genes from all four *lys3* mutants were examined and all contained defects likely to be deleterious (Supplementary Table S1 and Fig. 1D). The sequences of three of the four *lys3 PBF* alleles are disrupted by SNPs. Risø1508 and M1460 both have the same SNP (A173T) which causes an amino acid substitution (Q58L) in the DOF (<u>D</u>NA binding with <u>o</u>ne zinc <u>f</u>inger) DNA binding domain, consistent with Moehs *et al*. (2019). The glutamine residue, Q58 is highly conserved amongst orthologous proteins (Ensembl plants), with 27 out of 28 sequences examined having glutamine in this position. Risø19 has a single nucleotide polymorphism (G109T) which results in a nonsense mutation (E to STOP), which would prevent production of a full-length protein. In Risø18, various lines of evidence suggest that the entire *PBF* gene together with several additional genes on either side are deleted. Firstly, analysis of the chromosome 5H sequence data for Risø18 showed that sequence coverage around *PBF* was lacking (data not shown). Secondly, *PBF* could not be amplified by PCR with Risø18 DNA as template (Supplementary Figure S1). The presence of a large deletion around *LYS3* may also explain why we were unable to find any recombination close to this locus in the mapping lines derived from Risø18 (see 3.1).

### 3.3 Wheat PBF mutant grains have large embryos

To confirm that *PBF* is the gene responsible for controlling embryo size, we selected wheat cv. Cadenza TILLING lines (Krasileva et al., 2017) each affected in one of the wheat *PBF* homoeologs (TraesCS5A02G155900, TraesCS5B02G154100 and TraesCS5D02G61000) (Supplementary Table S2D). For *PBF-A*, no nonsense (premature termination codon) mutations were available and so we selected two lines (Cadenza1533 and Cadenza1807) with missense mutations (F48Y and T46I, respectively) in the DOF domain that are likely to be deleterious (SIFT score = 0.00). For the B- and D-genome *PBF* genes, we selected lines with nonsense mutations (Cadenza0903, Cadenza0904, respectively). Two triple mutant lines, Line 1 and Line 2, were created that contained the same B- and D-genome nonsense mutations but different A-genome missense mutations (Fig. 3A). Lines 2A and 2B contain the same three *PBF* mutations but were derived from different F_1_ grains.

**Figure 3.**
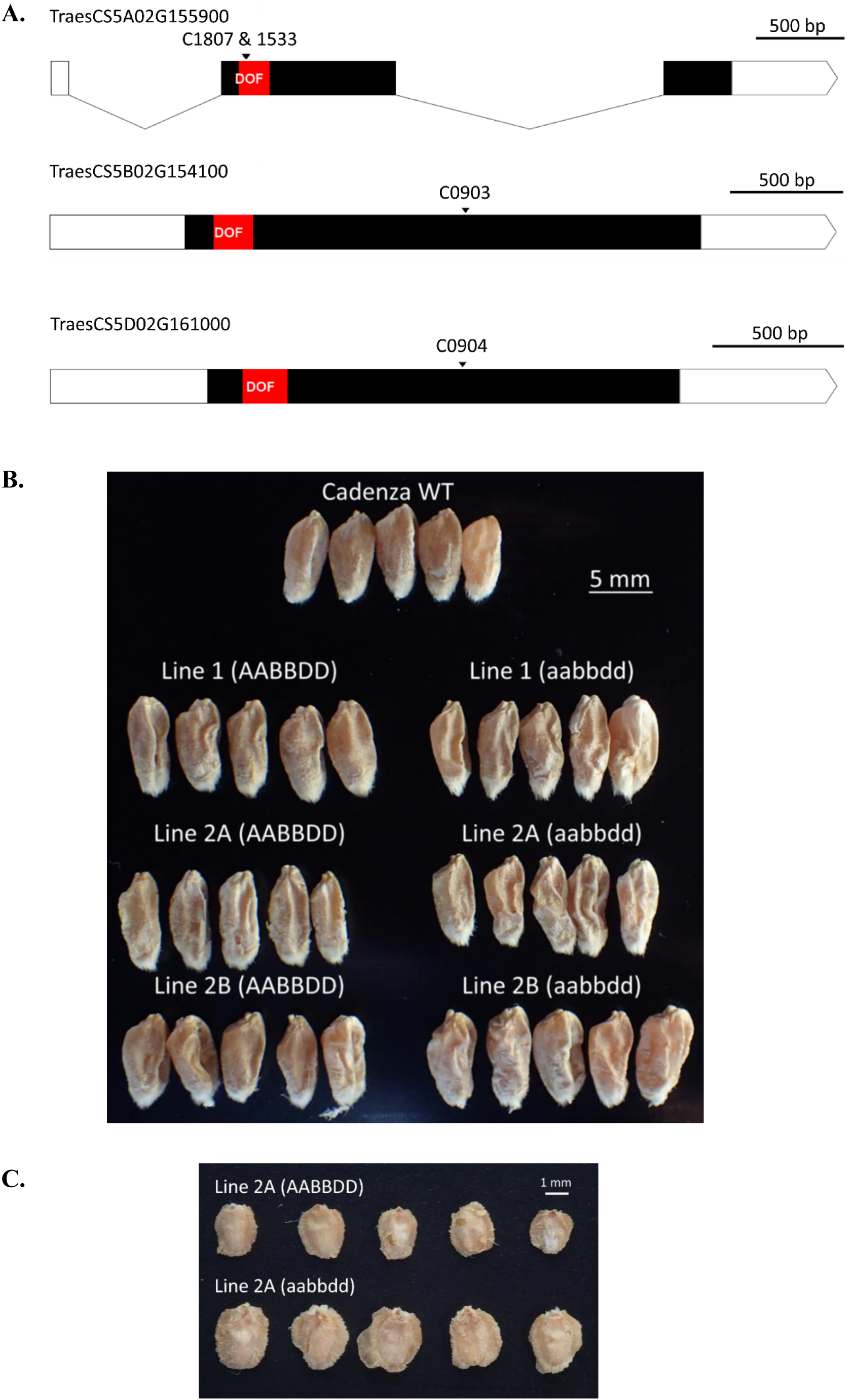
Wheat *PBF TILLING* mutants. Two wheat *PBF* triple mutant (aabbdd) lines were created by combining single A, B and D-homoeolog TILLING mutations in a single plant. Line 1 = Cadenza1533 x Cadenza0903 x Cadenza0904 (A missense x B nonsense x D nonsense). Line 2 = Cadenza1807 x Cadenza0903 x Cadenza0904 (A missense x B nonsense x D nonsense). Line 2A and 2B were derived from different F_1_ grains. A. A diagram of the *PBF* A, B and D gene structures. The exons are shown as bars and intron as lines. The coding regions are black and the DNA-binding DOF domain is red. The positions of the mutations are indicated by arrows. The two missense mutations are separated by 5 bp. C = Cadenza, NS = nonsense, MS = missense. B. Representative wild type (AABBDD) and mutant (aabbdd) F_3_ grains, and grains of the non-mutant, parental cultivar Cadenza are shown. All five grains of each type are from a single plant. All F_2_ plants were grown as a single batch. C. Representative wild type (AABBDD) and mutant (aabbdd) embryos are shown. Embryos are from F_3_ grains, as shown in Figure 3B.

Grains and embryos from the triple mutant plants, wild-type sibling plants and Cadenza were compared (Fig. 2B and Fig. 3). This showed that the grains of all TILLING lines (both wild-type and mutant) were more shrivelled than those of the non-mutant parent, Cadenza grown at the same time (Fig. 2B). We assume that the shrivelled appearance of the wild-type TILLING lines is due to mutations other than those in *PBF*.

The Line 1 triple mutant grains and embryos were indistinguishable from wild-type sibling controls in appearance and weight (Fig. 2B and Fig. 3). We assume that the A-genome missense mutant (F48Y) used to create Line 1 (Cadenza1533) had no deleterious impact on PBF functionality. In contrast, both Lines 2A and 2B (using Cadenza1807 as the A genome missense mutant, T46I) had increased absolute and relative embryo weights compared to their wild type sibling controls and the grains were more shrivelled than their wild-type sibling controls (Figs. 2B and Fig. 3). Thus, in wheat as in barley, PBF suppresses embryo growth.

### 3.4. Patterns of expression of PBF in barley

The Barley eFB Browser (www.bar.utoronto.ca/efp_barley; Druka *et al*., 2006; Winter *et al*., 2007) showed that *PBF* (HORVU5Hr1G048700) in cv. Morex is expressed in the developing grains (caryopsis) (Fig. 4A). Data for the developing embryo are available for one time point only: 22 DAF. *PBF* is expressed in the embryo at this time point, but at a low level compared with its expression in the rest of the caryopsis. *PBF* expression in other tissue types is very low or absent. A similar pattern of expression was seen for wheat cv. Azhurnaya. The wheat eFB Browser (www.bar.utoronto.ca/efp_wheat; Ramírez-González *et al.,* 2018; Winter *et al*., 2007) showed that expression for all three homoeologs was strong in the endosperm and that, for the one stage of development for which there is data, there was no expression in the embryo.

**Figure 4.**
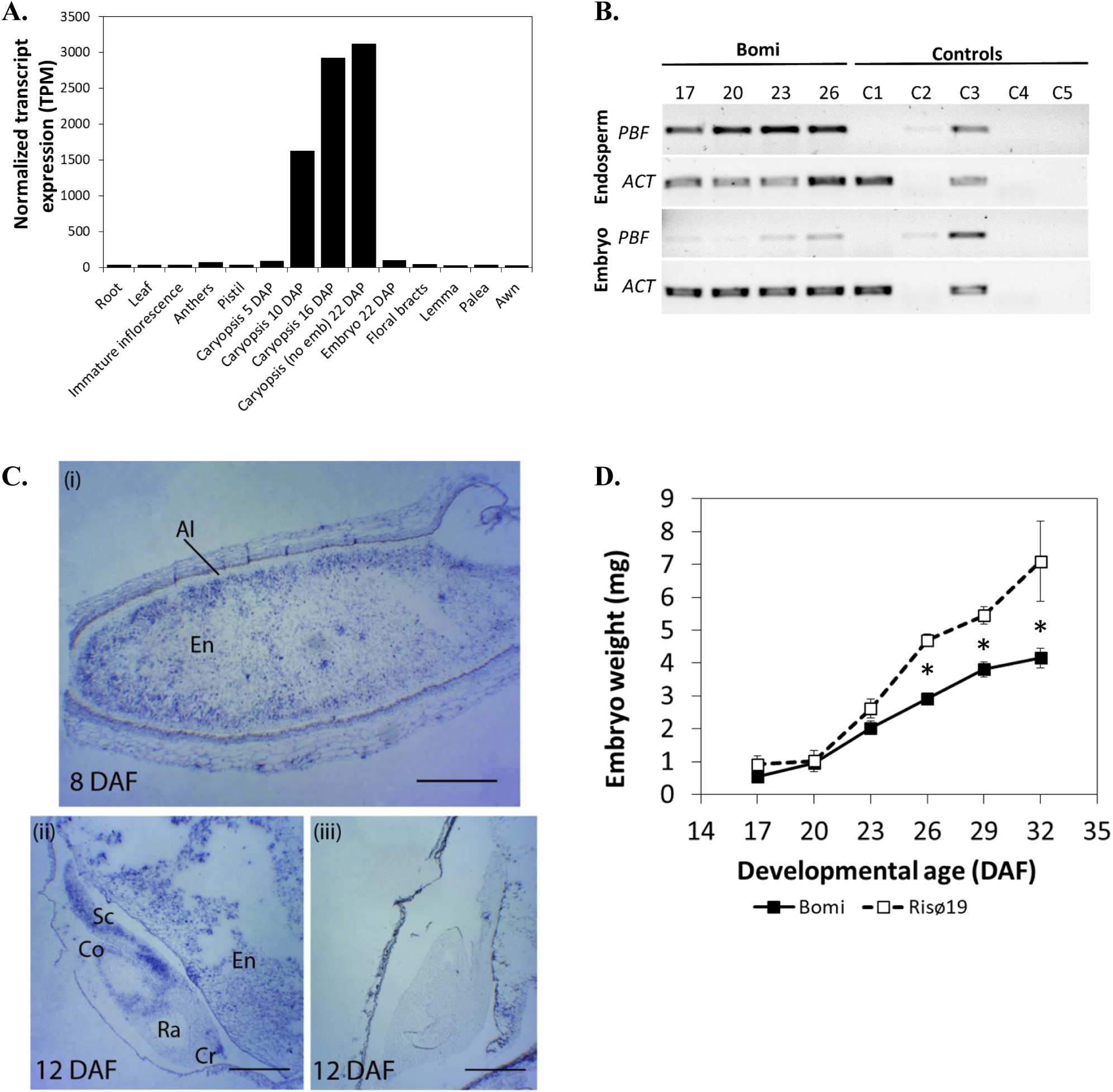
Patterns of expression of *PBF/LYS3* in barley. A. Tissue-specific expression according to the Barley eFP Browser 2.0 (www.bar.utoronto). Samples are from barley cv. Morex. Caryopsis (no Emb) means caryopsis without embryo. B. Temporal and tissue-specific patterns of *LYS3*/*PBF* expression in developing grains of wild type (Bomi) assessed by RT–PCR. Each cDNA sample was from a pool of 10-75 tissue samples each from an individual grain. Samples were collected from at least five spikes, each from a different plant. Amplicons were visualized on 1% agarose gels stained with SYBR Safe (Invitrogen, UK). Numbers above panels are DAF. *PBF* = *Prolamin Binding Factor* (HORVU5Hr1G048700). *ACT* = *ACTIN*.(HORVU1Hr1G047440), a constitutively expressed gene. Control reactions varied from the test reactions as follows. C1: contained cDNA from the barley *PBF* deletion mutant Risø18 at 26 DAF. No product was observed showing that the primers used were specific for *PBF.* C2: no DNase, no reverse transcriptase (RT). C3: no DNase. C4: no RT. C5: contained water used for PCR instead of cDNA. C. *In situ* localization of *PBF* in developing wild type (Bomi) grains at 8 and 12 DAF. (**i**) Longitudinal section of 8 DAF grain stained with antisense probe. *PBF* is expressed mostly in the starchy endosperm (En) cells. (**ii**) Longitudinal section of 12 DAF grain (including the embryo) stained with an antisense probe. *PBF* is expressed in the scutellum (Sc), coleoptile (Co) and coleorhiza tip (Cr). *PBF* is not expressed in the radicle (Ra). (**iii**) Longitudinal section of 12 DAF as in (ii) but stained with a sense probe (negative control). Scale bars are 0.5 mm (i) and 20 mm (ii and iii). D. Embryo fresh weight in developing grains of wild type (Bomi) and *lys3* mutant Risø19. Embryos were extracted from developing grains and immediately weighed. Values are means ± SE (n > 4) for 10 embryos extracted from the middle of at least three spikes. Values are significantly different (p<0.05, Tukey’s HSD after one-way Anova, mutant compared to wild type) for 26, 29 and 32 DAF (as indicated by asterisk). Error bars are included for all data and if not visible are smaller than the marker.

To compare with the data available for developing Morex embryos, and to expand upon it, the pattern of expression of *PBF* in developing embryos of the *LYS3* wild type, Bomi was determined using RT-PCR (Fig. 4B). Expression of *PBF* was detected in embryos at 23 and 26 DAF but the expression at 17 and 20 DAF was low and comparable to that seen in some of the control reactions suggesting that it is expressed at higher levels after 20 DAF. In comparison, *PBF* expression was detected in the endosperm at all stages of development tested. To investigate the temporal and tissue specific patterns of expression of *PBF* in young developing barley grains of cv Bomi further, we used *in situ* hybridization (Fig. 4C). This showed that at 8 DAF, *PBF* is expressed intensely in starchy endosperm cells and sparsely in aleurone cells. In the embryo at 12 DAF, *PBF* is expressed in the scutellum, the coleoptile and the tip of the coleorhiza.

To determine when during grain development the suppression of embryo growth occurs, we compared the fresh weights of developing Bomi embryos with those of the *lys3* mutant Risø19 (Fig. 4D). This showed that *lys3* mutant embryos were significantly larger than wild type embryos only after 23 DAF (Student’s *t*-test, p= 0.01). Thus, in wild-type embryos, the onset of *PBF* expression precedes the suppression of embryo growth.

### 3.5. In silico prediction of the downstream targets of wheat PBF

To understand how mutations in *PBF* effect both endosperm and embryo development and how these might be manipulated individually, we investigated the targets of this transcription factor. PBF is known to activate or repress the transcription of a number of genes by binding to specific sequences in their promotor regions called prolamin (TGTAAAG or CTTTACT) or pyrimidine (CCTTTT or AAAAGG) boxes (Mena *et al*., 1998; Mena *et al*., 2002). In developing barley endosperm, PBF target genes include B-hordein *(Hor2)* and trypsin-inhibitor BTI-CMe *(Itr1)* and in germinating barley grains, cathepsin B-type protease *(Al21)* and α-Amylase *(Amy2/32b)* are PBF targets (Mena *et al*., 1998; Diaz *et al*., 2002; Mena *et al*., 2002). Nothing is yet known about PBF targets in developing barley embryos.

At present, the data and tools available for target gene analysis are more advanced for wheat than for barley. Wheat transcription-factor targets can be predicted using a GENIE3 network, which was created using 850 diverse RNAseq samples from wheat (Ramírez-González et al, 2018). Using the GENIE3 network to predict the downstream targets of the wheat *PBF* homoeologs, we identified >450 gene targets. We found that these target genes were enriched for a wide range of gene ontology (GO) annotations (Supplementary Fig. S2 and Fig. 5) but the two categories that were the most significantly enriched were organ development and starch metabolism.

**Figure 5.**
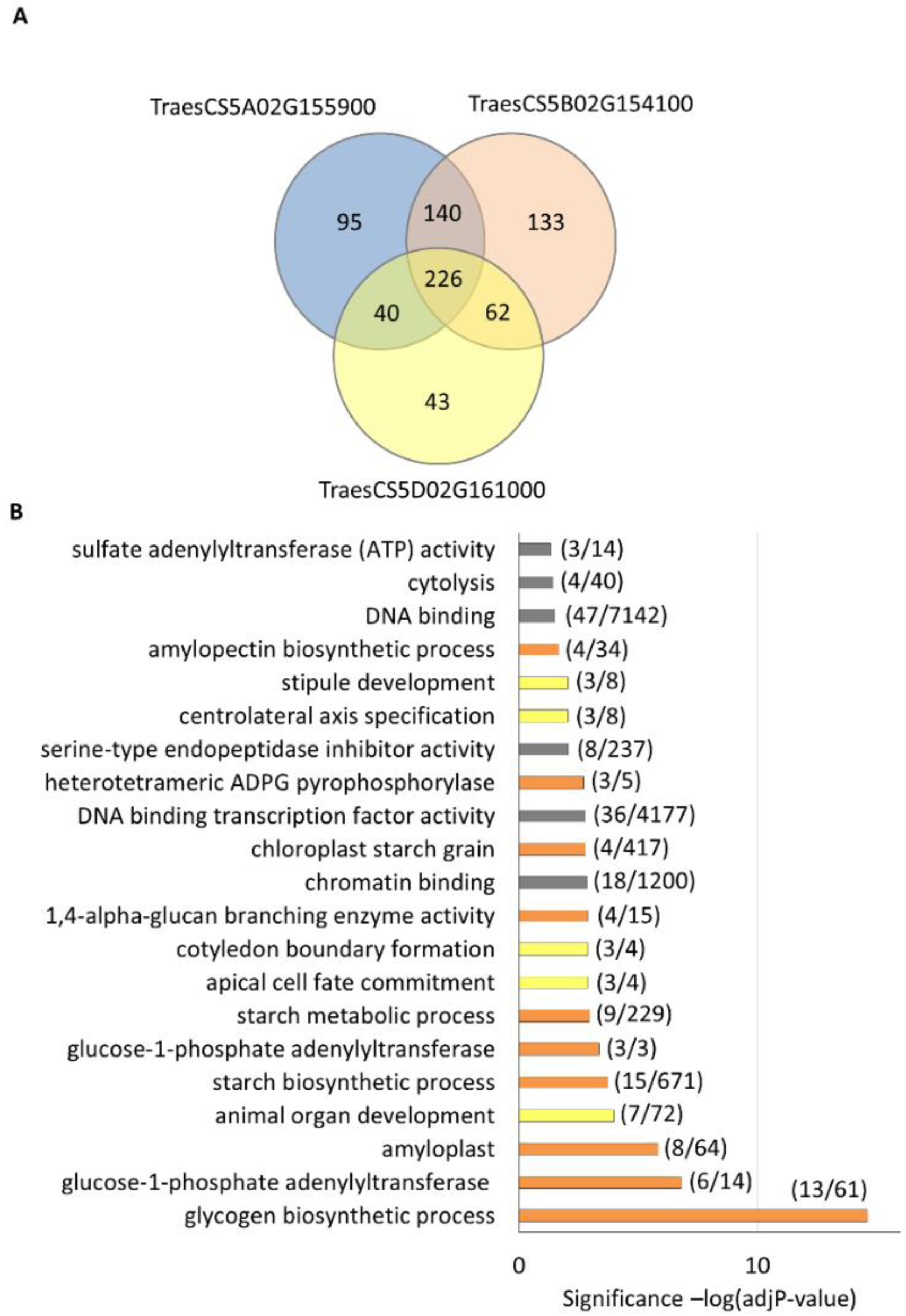
Downstream targets of wheat *PBF*. The downstream target genes of *PBF* were identified using a GENIE3 network constructed using 850 RNA-seq samples (Ramírez-González et al., 2018). A. A comparison of the target genes of each of the wheat *PBF* homoeologs. The number of predicted target genes common to all three homoeologs is 226. B. The 226 common target genes were analysed for evidence of enrichment with respect to molecular function. The top-ranking functional categories (adjusted p value <0.05) are presented with the significance of the enrichment [−log(adjP-value)]. In brackets are the numbers of genes detected in each category with respect to all GO terms annotated so far for wheat RefSeqv1 gene models (Ramírez-González *et al*., 2018). GO terms related to starch metabolism are in orange, organ development in yellow, and others in grey.

To investigate whether PBF directly regulates genes known to be involved in controlling embryo size, we looked at the downstream targets of PBF for homologues of three genes known to give rise to large embryo/small endosperm mutant phenotypes in rice (*Oryza sativa* L.). The genes examined were *GIANT EMBRYO* (*GE*), encoding a cytochrome P450 (CYP78A13) (Satoh and Omura, 1981; Nagasawa *et al*., 2013); *BIGE1*, encoding a MATE (Multidrug-And-Toxic-compound-Extrusion) type transporter protein (Suzuki *et al*., 2015); and *LARGE EMBRYO*, encoding a C3HC4-type RING (Really Interesting New Gene) finger protein of unknown function (Lee *et al*., 2019). None of these three genes was a predicted target of the wheat *PBF* ortholog.

To determine whether *PBF* participates in the same regulatory networks (or modules) as any of the other large-embryo genes (*GE*, *BIGE1* and *LARGE EMBRYO*), we used co-expression networks built using WGCNA (Ramírez-González *et al.,* 2018) (Supplementary Table S3). For the grain-specific network, the wheat homoeologs of *PBF* are in module 13, *BIGE1* in module 2 and *GE* in module 2 (B genome) or 9 (A and D genome). The *LARGE EMBRYO* homoeologs are in module 0, suggesting that the expression of these genes is invariant across all grain samples. Thus, for wheat it is likely that *PBF* regulates embryo size independently of the other three genes whilst *BIGE1* and the B genome of *GE* both operate within the same regulatory network.

PBF was so named because it was found to recognise a conserved cis-element, the prolamin box, in the promotor regions of prolamin seed storage proteins (Wu and Messing, 2012). In barley, PBF was also shown to bind to the pyrimidine box of genes expressed in aleurone cells during seed germination (Mena et al., 2002). To investigate whether PBF regulates genes involved in starch metabolism by binding to prolamin or pyrimidine boxes in their promotor regions, as is the case for other genes regulated by PBF, we analysed the promotor regions of 27 genes involved in starch synthesis in the endosperm (Supplementary Table S4). This showed that ten of the starch genes have a prolamin box within the 2-kbp upstream regulatory (promotor) region of at least one homoeologue, all 27 genes have a pyrimidine box and 11 genes were predicted by GENIE3 to be downstream targets of PBF. Ten starch genes have both prolamin and pyrimidine boxes of which only four were predicted targets. Seven additional genes were predicted targets of PBF and these lacked a prolamin box. Thus, for this limited data set, there is little or no correlation between the presence of the two known PBF regulatory sequences in starch gene promotors and their interactions with PBF as predicted from gene expression studies. Despite its name, PBF, like other transcription factors, probably interacts with a range of DNA regulatory sequences.

## 3. Discussion

We showed by genetic mapping that the *LYS3* gene encodes a transcription factor called Prolamin Binding Factor (PBF), and that one of the many effects of mutations in this gene is an increase in embryo size. All four *lys3* alleles have deleterious mutations in the *PBF* gene (Supplementary Table S1) and wheat TILLING mutants with mutations in *PBF* also have enlarged embryos (Fig. 3), confirming a role for *PBF* in determining organ size in Triticeae grains. The *LYS3* gene was also identified as *PBF* independently by Moehs *et al*. (2019). However, they studied the effect of *LYS3* on hordein content in one mutant, *lys3a* and they did not measure embryo size.

The severity of the effects of *lys3* alleles on embryo size varies, with Risø1508 and M1460 having larger embryos than either Risø18 or Risø19 (Fig. 1A). The reasons for this are currently unknown but the two least-severe mutants, Risø18 or Risø19 both lack the PBF protein (due to nonsense mutations), whilst the two mutants with the largest embryos, Risø1508 and M1460 have the same missense mutation (A173T) affecting their DOF domains. Proteins encoded by genes with missense mutations can be partially defective, rather than entirely absent or entirely defective. It is therefore possible that the missense mutation in Risø1508 and M1460 weakens or eliminates the ability of PBF to bind to DNA, but does not affect its interactions with other regulatory proteins. This hypothesis requires investigation.

Previous studies have shown that *PBF* is expressed in starchy endosperm and aleurone cells during barley grain development (Mena *et al*., 1998; Mena *et al*., 2002) and in the aleurone during germination (Mena *et al*., 2002) but no expression in developing embryos was detected using Northern blots (Mena *et al*., 2002). In this study, we confirmed the expression of *PBF* in starchy endosperm and aleurone cells in developing barley grains. We also showed using RT-PCR and *in situ* hybridization that *PBF* is expressed in developing embryos (Fig. 4B, C). *PBF* expression in the embryo increased during grain development but expression was detected prior to the time point when embryo growth in wild-type and mutant grains diverged. These data suggest that control of embryo growth could be mediated by *PBF* expression in the embryo and not indirectly via its expression in the endosperm. Thus, gene regulation by PBF in embryos may involve a regulatory pathway that is independent from the PBF pathway operating in the endosperm.

To understand the regulatory pathways downstream of PBF in developing Triticeae grains, we used *in silico* prediction of target genes based on wheat RNAseq data (Ramírez-González et al 2018; Harrington et al 2019). This indicated that PBF in developing grains is involved in the regulation of a wide range of processes but the two most significant categories were organ development and starch metabolism (mirroring its principal roles in embryo growth and endosperm starch synthesis, respectively). Other genes known to cause large embryos when mutated in rice were considered as potential downstream targets of PBF but no such interactions were predicted from analysis of published RNAseq data. We also looked for evidence of co-expression of *PBF* with these other genes in wheat grains. No such evidence was found. Thus, the PBF transcriptional network that controls embryo growth remains unknown. However, it must be noted that the expression data used for this *in silico* analysis included three embryo samples only (Ramírez-González et al 2018). More RNAseq data will need to be gathered for developing embryos to test these ideas further.

In maize, as in wheat and barley, there is a *PBF* gene that regulates starch metabolism suggesting that the regulation of endosperm starch synthesis by *PBF*s may be a common feature amongst grasses. Maize PBF controls the expression of four starch biosynthetic genes, *Sh2* and *Bt2* that encode, respectively, the cytosolic large and small subunits of ADP glucose pyrophosphorylase (AGPase); *SBEI* encoding starch branching enzyme I and *Su1* that encodes isoamylase 1. The expression of both AGPase subunit genes was down-regulated in maize *PBF* RNAi lines whilst the other two genes were up-regulated (Zhang *et al.,* 2016). Orthologs of all four genes in wheat are also predicted target genes of wheat *PBF* (Supplementary Table S4) suggesting that the *PBF* transcription factors in maize and the Triticeae species interact with similar sets of downstream genes. However, there is no information to date to suggest that *PBF* controls embryo size in maize.

Large-embryo size in cereals is associated with increased nutritional value because embryos are rich in protein, vitamins, oil and non-starch carbohydrates. There is also evidence of human health benefits from eating large-embryo rice grain products (Zhang *et al.,* 2005; Lee *et al.,* 2016; Jung *et al.,* 2017). As wholegrain products are becoming increasingly popular for human consumption, cereal grains with large embryos offer new opportunities for cereal grain improvement. We have shown that barley and wheat with mutations in *PBF* have enlarged embryos but unfortunately, they also have reduced grain size due to endosperm defects. However, the possibility that the regulatory networks controlled by PBF in the endosperm and embryo are independent provides an opportunity to explore their independent manipulation. Further work is in progress to pursue this idea.

## Acknowledgements

This work was supported by the Biotechnology and Biological Sciences Research Council (BBSRC), UK [grant numbers BB/L023156/1 and BB/P016855/1] and an Anniversary Future Leader Fellowship BB/M014045/1 to PB. CUS thanks TETFUND Nigeria for a PhD training grant. JD and MK were supported by ERDF project “Plants as a tool for sustainable global development” (No. CZ.02.1.01/0.0/0.0/16_019/0000827). We thank Zdeňka Dubská, Romana Šperková, Jan Vrána and Jitka Weiserová for sample preparation, chromosome sorting and DNA amplification.

## AUTHOR CONTRIBUTIONS

KT designed the project, with advice from CU, PB and BBHW. NS generated and analysed the mapping populations. MK managed chromosome sorting and performed purity checks under the supervision of JD. Bioinformatic analysis of the sequenced chromosome was done by BS with help and advice from CU and BBHW. BOL carried out the bulk of the other experimental work on barley and BOL and PB carried out the wheat *PBF* target gene predictions. TC and MC were responsible for the production and analysis of the wheat TILLING mutants. CUS performed the in situ analysis, and SD supervised this work. BOL and KT wrote the paper together and all authors proof-read the paper and provided feedback. Authors, apart from the first and last, are listed alphabetically.

---

**Supplementary Figure S1.**
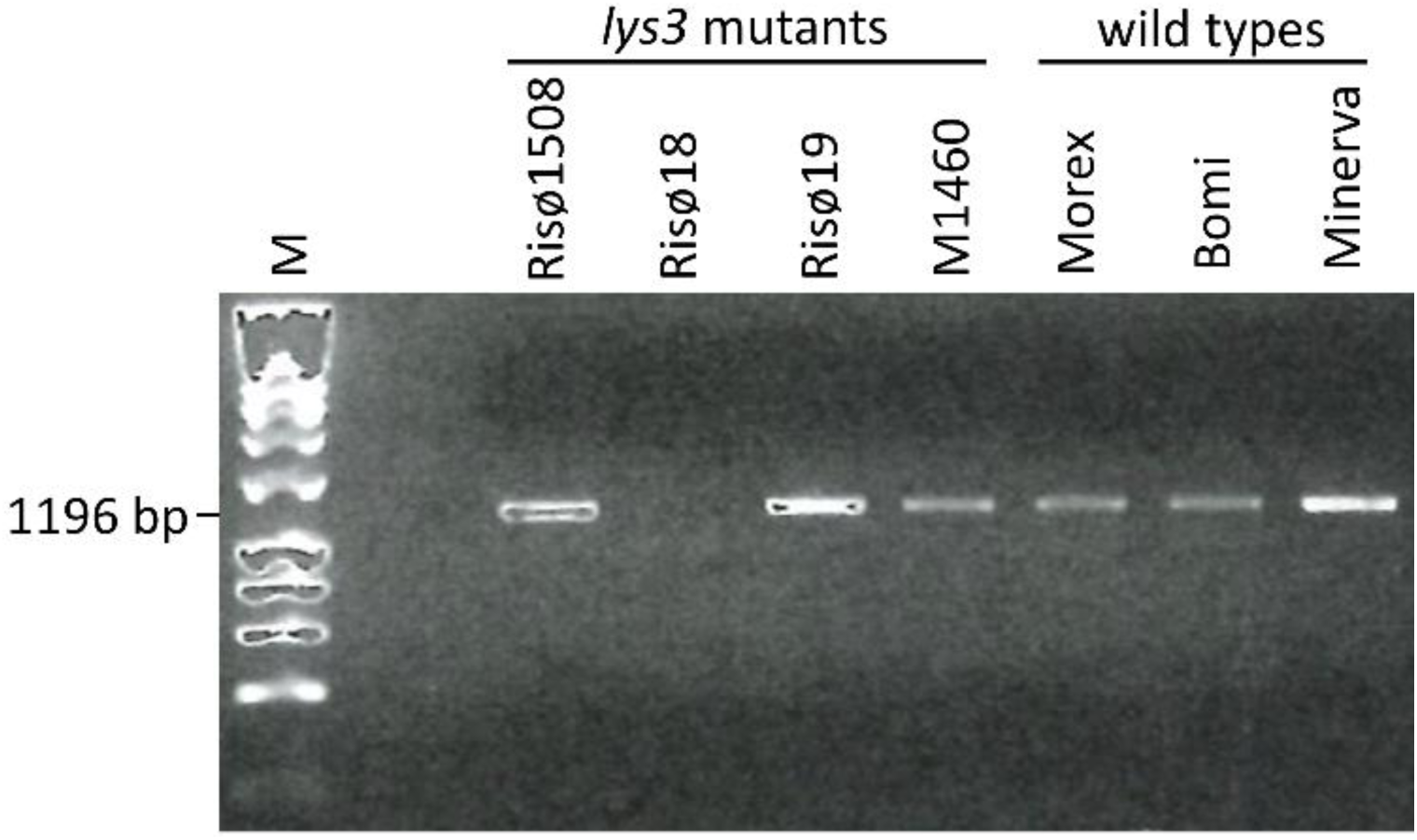
Comparison of the *PBF* alleles in all four *lys3* mutant lines. Barley *PBF* genomic sequences were amplified by PCR and amplicons were separated by electrophoresis. M = molecular weight ladder (Bioline Ladder 1 kbp). The size of the wild type *PBF* amplicon is indicated.

**Supplementary Figure S2.**
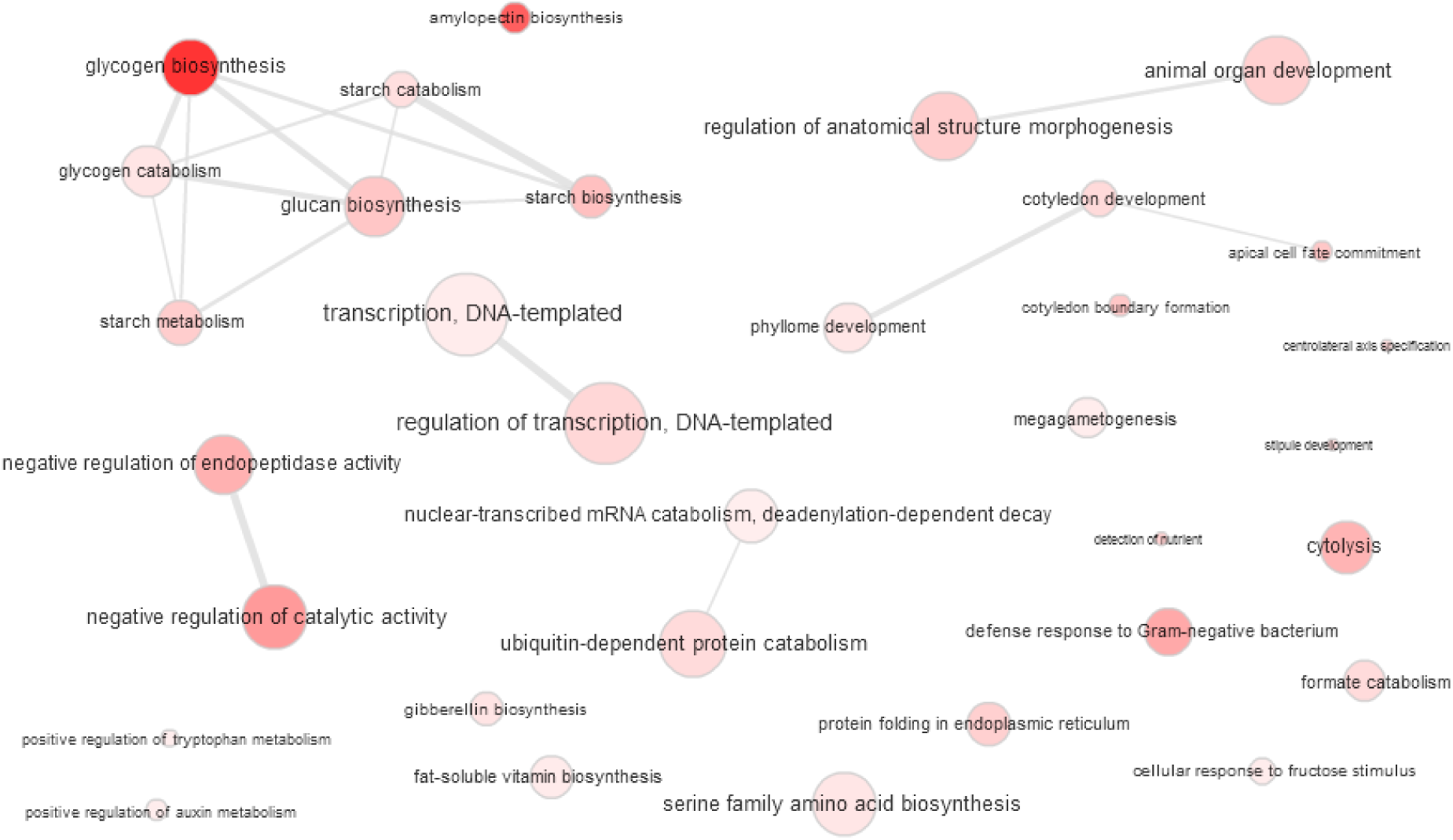
Predicted downstream targets of *PBF* in wheat. GENIE3 analysis of 850 wheat RNASeq samples revealed putative downstream targets of *PBF* (Fig. S2). The gene ontologies (GOs) were reduced and visualized using enrichment analysis (http://revigo.irb.hr/revigo.jsp; Ramírez-González *et al*., 2018). Bubble colour indicates p-value and bubble size the frequency of the GO term in the GO database. Highly similar GO terms are linked by edges in the graph.

**Supplementary Table S1.** The *lys3* mutants of barley. Comparison of the four known *lys3* mutant lines: their origins and the effects of the mutations on *PBF* gene and protein. See also Fig. 1D.

| Mutant | Wild type | Mutagen | Original selection criteria | Reference | Change to cDNA | position | Effect | Change to protein |
| --- | --- | --- | --- | --- | --- | --- | --- | --- |
| Risø18 | Bomi | Na azide | Unknown | Munck *et al.*, 1992 | - | - | Deletion | - |
| Risø19 | Bomi | Na azide | Unknown | Munck *et al.*, 1993 | G/T | 109 | Nonsense/Stop | E/* |
| Risø1508 | Bomi | Ethylenimine | High lysine | Tallberg *et al.*, 1973 | A/T | 173 | Missense | Q/L |
| M1460 | Minerva | Na azide | Low β-glucan | Aastrup *et al.*, 1983 | A/T | 173 | Missense | Q/L |

**Supplementary Table S2.**
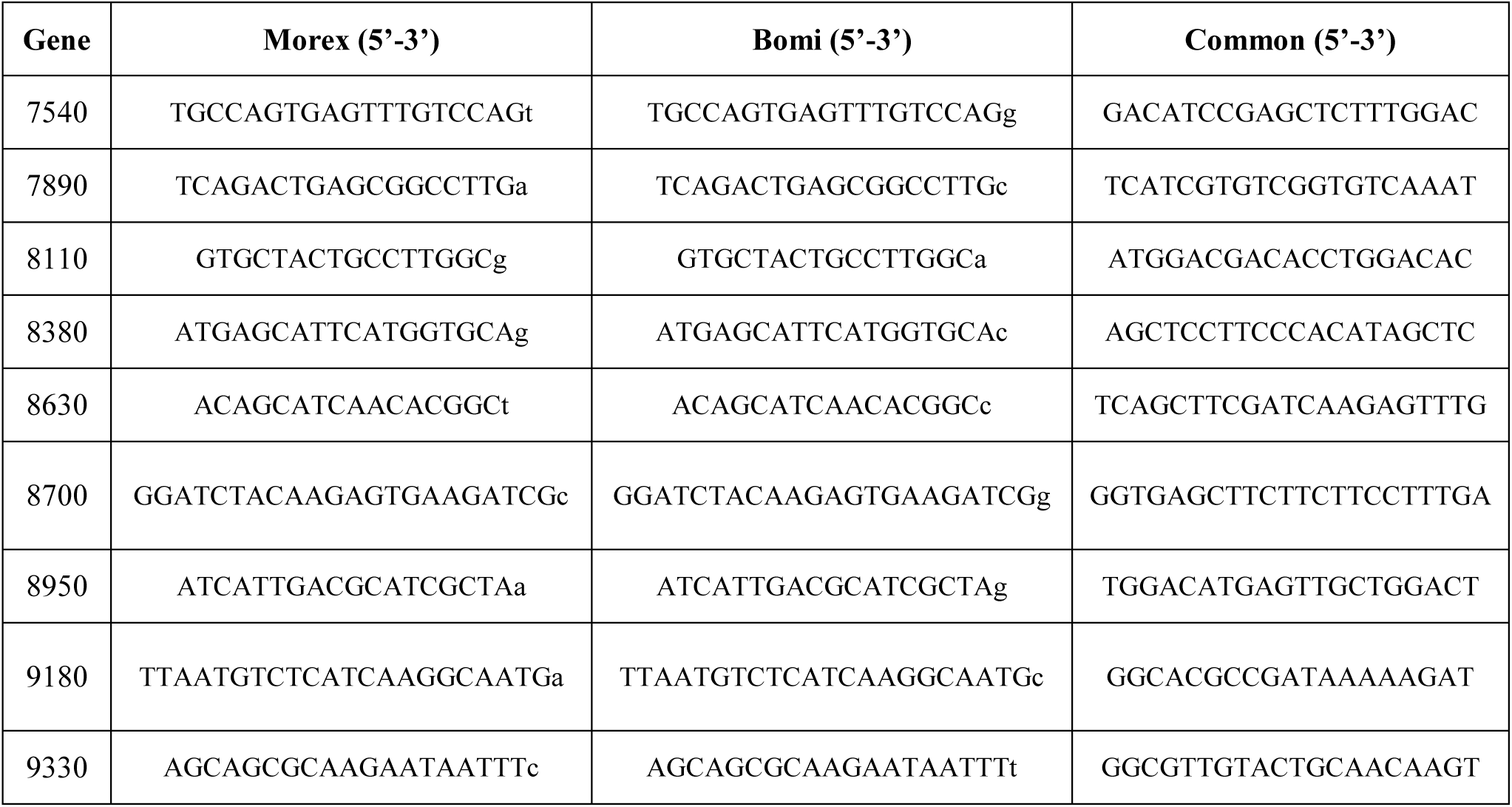
Primers and assay conditions. A. KASP primers for barley genotyping. Primers were preceded by the VIC sequence (GAAGGTCGGAGTCAACGGATT) (Morex and *LYS3*) or by FAM sequence (GAAGGTGACCAAGTTCATGCT) (Bomi and *lys3*). The discriminatory bases are small case. The ratio of FAM/VIC/Common primers used was 12/12/30, except for the genotyping primers for HORVU5Hr1G048380, which were in the ratio 12/6/30. The KASP reaction (in a final volume of 5 μl) contained: 2.5 μl 2xKASP reagent, 0.07 μl of primer mix and 2.5 μl of gDNA template (1 ng/μl). The PCR conditions were: 94 °C for 15 min, 10 cycles of (94 °C for 20 s, 65 °C for 1 min and -0.8 °C/cycle to 57 °C), 10 cycles of (94 °C for 20 s, 57 °C for 1 min) and then reprise 5 to 10 cycles of (94 °C for 20 s, 57 °C for 1 min). Abbreviated gene names (xxxx) are as in Table 1 (i.e. HORVU5Hr1G04xxxx).

**Supplementary Table S2.**
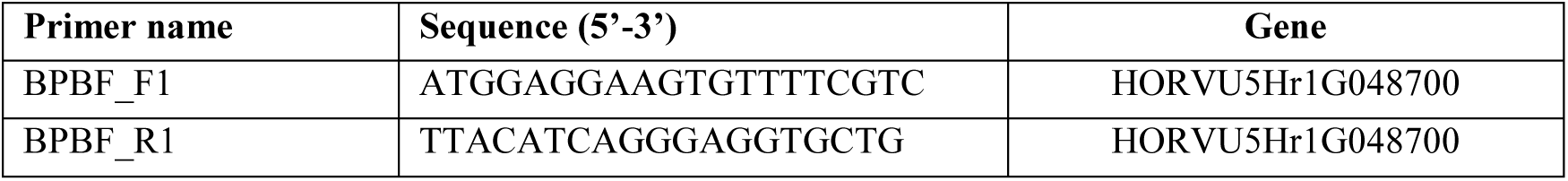
B. Primers used to amplify barley *PBF* genomic sequence. The PCR reaction (in a final volume of 20 μl) contained: 1 μl of gDNA template (2 ng), 0.5 μl of 10 mM forward primer, 0.5 μl of 10 mM reverse primer, 0.6 μl of 10 mM dNTPs, 2 μl of 10 x PCR buffer, 0.2 μl of Phusion High-Fidelity DNA Polymerase (Thermo Fisher Scientific, UK) and water. The PCR conditions were: 98 °C for 30 s, 35 cycles of (98 °C for 10 s, 58 °C for 60 s and 72 °C for 20 s) and then 72 °C for 5 min.

**Supplementary Table S2.**
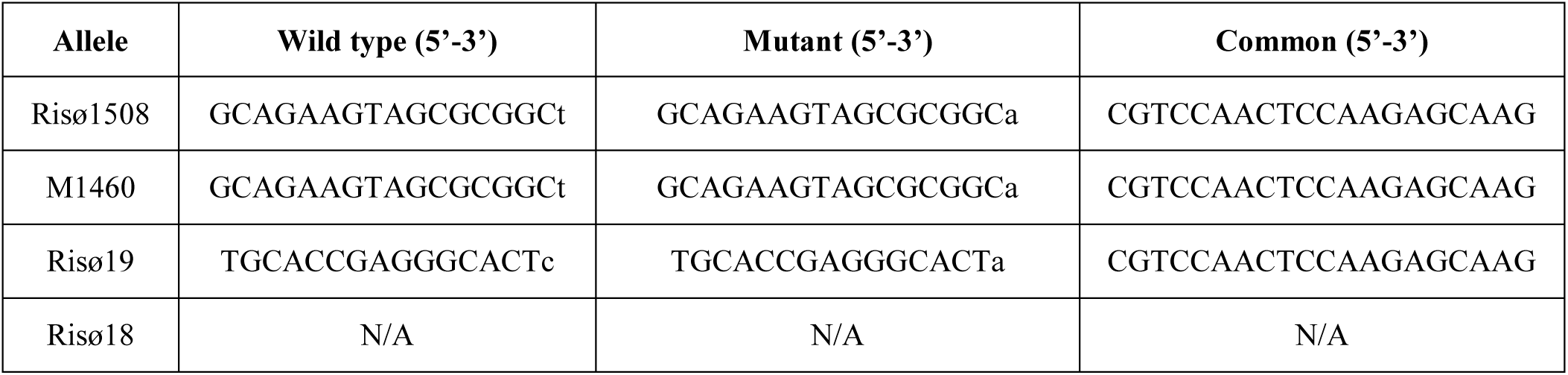
C. Barley *PBF* (HORVU5Hr1G048700) allele-specific KASP primers. Primers were preceded by the VIC sequence (GAAGGTCGGAGTCAACGGATT) (*lys3*) or by FAM sequence (GAAGGTGACCAAGTTCATGCT) (Bomi). The discriminatory bases are small case. The ratio of FAM/VIC/Common primers used was 12/12/30. The PCR reactions and conditions were as for (A). Note that no primers are given for Risø18 because the genomic region containing this allele is deleted.

**Supplementary Table S2.**
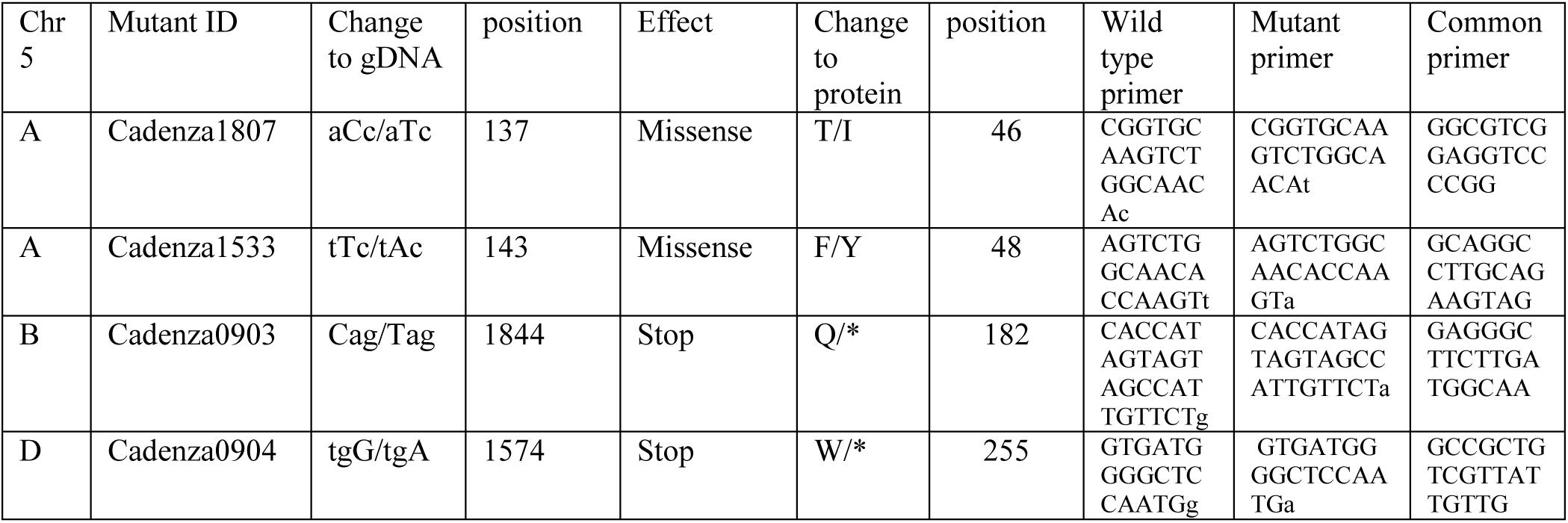
D. Wheat TILLING mutants and KASP primers. Primers were designed for KASP assays and were preceded by the VIC sequence (GAAGGTCGGAGTCAACGGATT) (wild-type) or by FAM sequence (GAAGGTGACCAAGTTCATGCT) (mutant). The discriminatory bases are small case. The ratio of FAM/VIC/Common primers used was 12:12:15 (Cadenza1807), 6:12:30 (Cadenza1533) or 12:12:30 (Cadenza0903 and Cadenza0904). The KASP reaction was carried out as described by the manufacturer (https://www.lgcgroup.com).

**Supplementary Table S2.**
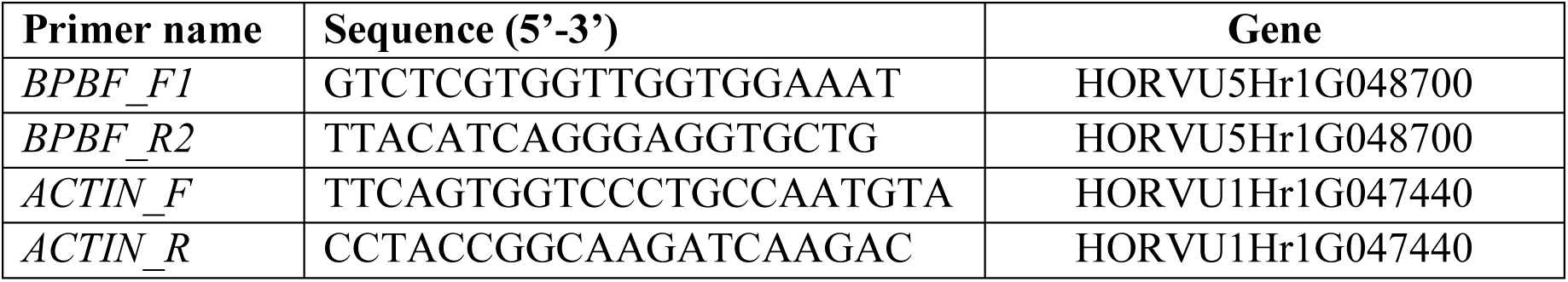
E. Primers for analysis of gene expression by RT-PCR. The RT-PCR reaction (in a final volume of 20 μl) contained: 1 μl of cDNA template, 0.5 μl of 10 mM forward primer, 0.5 μl of 10 mM reverse primer, 0.6 μl of 10 mM dNTPs, 2 μl of 10 x PCR buffer, 0.2 μl of FastStart *Taq* DNA polymerase (Roche, 04659163103) and water. The PCR conditions were: 95 °C for 3 min, 35 cycles of (95 °C for 30 s, 58 °C for 30 s and 72 °C for 30 s or 1 min) and then 72 °C for 5 min. Reactions were performed in triplicate and *ACTIN*, a constitutively-expressed gene, was used as a control.

**Supplementary Table S2.**
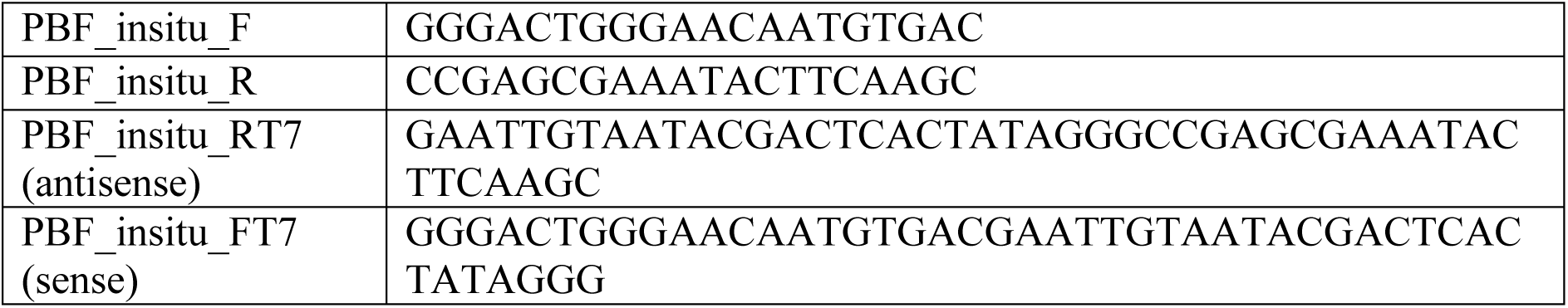
F. Primers for *in situ* expression analysis. *In situ* hybridization was as described previously (Drea, *et al*., 2005; Opanowicz *et al.,* 2010).

**Supplementary Table S3.** WGCNA co-expression analysis. Analysis of co-expression of *PBF* and other large-embryo genes (*GE*, *BIGE1* and *LARGE EMBRYO*) (Ramírez-González *et al.,* 2018). The co-expression Module (network) associated with each gene for four different tissues is indicated. NA = not analysed (because the gene is not expressed in this tissue and so was not included in the network).

| Gene type | Gene ID | Module |  |  |  |
| --- | --- | --- | --- | --- | --- |
|  |  | Grain | Leaf | Root | Spike |
| *GE* | TraesCS2A01G175700 | 9 | 0 | 11 | 1 |
|  | TraesCS2B01G201900 | 2 | 1 | 18 | 1 |
|  | TraesCS2D01G183000 | 9 | 1 | 18 | 0 |
| *Lys3/PBF* | TraesCS5A01G155900 | 13 | NA | NA | NA |
|  | TraesCS5B01G154100 | 13 | NA | NA | NA |
|  | TraesCS5D01G161000 | 13 | NA | NA | NA |
| *LE* | TraesCS4A01G271200 | 0 | 0 | 2 | 0 |
|  | TraesCS4B01G042900 | 0 | 0 | 2 | 17 |
|  | TraesCS4D01G040200 | 0 | 0 | 2 | 0 |
| *BIGE1* | TraesCS4A01G350200 | 2 | 0 | 7 | 12 |
|  | TraesCS5B01G522900 | 2 | 0 | 1 | 12 |
|  | TraesCS5D01G521600 | 2 | 0 | 0 | 0 |

**Supplementary Table S4.** Analysis of the promotor regions of wheat starch-related genes. A set of wheat genes involved in starch synthesis was selected and their promotor regions (2 kb upstream sequences) were identified in the IWGSC RefSeqv1 wheat cv Chinese Spring (CS) genome sequence as part of the WGIN Wheat Promotome Capture project (http://www.wgin.org.uk/). The presence of *PBF* regulatory motifs (prolamin box, TGTAAAG or CTTTACT; pyrimidine box, CCTTTT or AAAAGG) in the promotor regions is indicated. Genes predicted by GENIE3 analysis to be down-stream targets of *lys3 PBF* are also indicated (see Fig. 5).

| Gene function or product | Gene designation | IWGSC gene ID | Homoeolog present in CS | Prolamin box present | Pyrimidine box present | Predicted target genes of *lys3 PBF* (GENIE3). |
| --- | --- | --- | --- | --- | --- | --- |
| **ADPG pyrophosphorylase large subunit** | *APL1/AGPL3* | TraesCS5A02G472000 | Yes | No | Yes | No |
|  |  | TraesCS5B02G484700 | Yes | No | Yes | No |
|  |  | TraesCS5D02G484500 | Yes | No | Yes | No |
| **ADPG pyrophosphorylase large subunit** | *APL3/AGPL1* | TraesCS1A02G419600 | Yes | Yes | Yes | Yes |
|  |  | TraesCS1B02G449700 | Yes | Yes | Yes | Yes |
|  |  | TraesCS1D02G427400 | Yes | Yes | Yes | Yes |
| **ADPG pyrophosphorylase small subunit** | *APS2/AGPS2* | TraesCS5A02G292900LC | Yes | No | Yes | No |
|  |  | TraesCS5B02G175800 | Yes | No | Yes | No |
|  |  | TraesCS5D02G182600 | Yes | No | No | No |
| **ADPG pyrophosphorylase small subunit** | *APS1/AGPS1/Bt2* | TraesCS7A02G287400 | Yes | Yes | Yes | Yes |
|  |  | TraesCS7B02G183300 | Yes | No | Yes | Yes |
|  |  | TraesCS7D02G284900 | Yes | No | Yes | Yes |
| **Disproportionating enzyme I** | *DPE1* | TraesCS2A02G159300 | Yes | No | Yes | No |
|  |  | TraesCS2B02G184900 | Yes | No | Yes | No |
|  |  | TraesCS2D02G166600 | Yes | No | Yes | No |
| **Disproportionating enzyme II** | *DPE2* | TraesCS2A02G123800 | Yes | No | Yes | No |
|  |  | TraesCS2B02G145700 | Yes | No | Yes | No |
|  |  | TraesCS2D02G126600 | Yes | No | Yes | No |
| Floury endosperm6 | *FLO6* | TraesCS4A02G284000 | Yes | No | No | No |
|  |  | TraesCS4B02G029700 | Yes | No | Yes | No |
|  |  | TraesCS4D02G026700 | Yes | No | Yes | No |
| Granule-bound starch synthase I | *GBSSI/waxy* | TraesCS7A02G070100 | Yes | No | Yes | Yes |
|  |  | TraesCS7D02G064300 | No | No | No | Yes |
|  |  |  | Yes | No | Yes | No |
| Granule-bound starch synthase II | *GBSSII* | TraesCS2A02G373600 | Yes | No | Yes | No |
|  |  | TraesCS2B02G390700 | Yes | Yes | Yes | No |
|  |  | TraesCS2D02G369800 | Yes | No | Yes | No |
| Isoamylase I | *ISA1* | TraesCS7A02G251400 | Yes | No | Yes | Yes |
|  |  | TraesCS7B02G139700 | Yes | No | Yes | Yes |
|  |  | TraesCS7D02G249500 | Yes | No | Yes | Yes |
| Isoamylase II | *ISA2* | TraesCS1A02G247100 | Yes | No | Yes | No |
|  |  | TraesCS1B02G257700 | Yes | Yes | Yes | No |
|  |  | TraesCS1D02G246300 | Yes | No | Yes | No |
| Isoamylase III | *ISA3* | TraesCS5A02G248700 | Yes | No | Yes | No |
|  |  | TraesCS5B02G246400 | Yes | No | Yes | No |
|  |  | TraesCS5D02G255800 | Yes | No | Yes | No |
| Limit dextrinase | *PUL* | TraesCS7A02G133500 | Yes | Yes | Yes | Yes |
|  |  | TraesCS7B02G034600 | Yes | No | Yes | Yes |
|  |  | TraesCS7D02G133100 | Yes | No | Yes | Yes |
| Maltose transporter | *MT* | TraesCS2A02G436000 | Yes | No | Yes | No |
|  |  | TraesCS2B02G457100 | Yes | No | Yes | No |
|  |  | TraesCS2D02G433900 | Yes | No | Yes | No |
| Plastidial ADPG transporter | *OsBT1-1* | TraesCS6A02G175100 | Yes | No | Yes | Yes |
|  |  | TraesCS6B02G210000 | Yes | No | Yes | Yes |
|  |  | TraesCS6D02G168200 | Yes | No | Yes | Yes |
| Starch branching enzyme I | *BEI* | TraesCS7A02G549300 | Yes | No | Yes | Yes |
|  |  | TraesCS7B02G472300 | Yes | No | Yes | Yes |
|  |  | TraesCS7D02G535600 | Yes | No | Yes | Yes |
| Starch branching enzyme IIa | *BEIIa* | TraesCS2A02G293400 | Yes | No | Yes | No |
|  |  | TraesCS2B02G309500 | Yes | No | Yes | No |
|  |  | TraesCS2D02G290800 | Yes | Yes | Yes | No |
| Starch branching enzyme IIb | *BEIIb/Ae* | TraesCS2A02G310300 | Yes | No | Yes | Yes |
|  |  | TraesCS2B02G327300 | Yes | No | Yes | Yes |
|  |  | TraesCS2D02G308600 | Yes | No | Yes | Yes |
| Starch phosphorylase (cytosol) | *PHOH* | TraesCS3A02G366300 | Yes | No | Yes | No |
|  |  | TraesCS3B02G397900 | Yes | No | Yes | No |
|  |  | TraesCS3D02G359200 | Yes | No | Yes | No |
| Starch phosphorylase (plastidial) | *PHOL* | TraesCS5A02G395200 | Yes | Yes | Yes | Yes |
|  |  | TraesCS5B02G400000 | Yes | No | Yes | Yes |
|  |  | TraesCS5D02G404500 | Yes | Yes | Yes | No |
| Starch synthase I | *SSI<sup>c</sup>* | TraesCS7A02G120300 | Yes | No | No | No |
|  |  | TraesCS7B02G018600 | Yes | No | No | No |
|  |  | TraesCS7D02G117800 | Yes | No | Yes | Yes |
| Starch synthase IIa | *SSIIa/SSII-3* | TraesCS7A02G189000 | Yes | No | Yes | No |
|  |  | TraesCS7B02G093800 | Yes | Yes | Yes | No |
|  |  | TraesCS7D02G190100 | Yes | Yes | No | No |
| Starch synthase IIb | *SSIIb/SSII-2* | TraesCS6A02G307800 | Yes | No | Yes | No |
|  |  | TraesCS6B02G336400 | Yes | No | Yes | No |
|  |  | TraesCS6D02G287000 | Yes | No | Yes | No |
| Starch synthase IIc | *SSIIc/SSII-1* | TraesCS1A02G137200 | Yes | No | Yes | No |
|  |  | TraesCS1B02G155700 | Yes | No | Yes | No |
|  |  | TraesCS1D02G138100 | Yes | No | Yes | No |
| Starch synthase IIIa | *SSIIIa/SSIII-2* | TraesCS1A02G091500 | Yes | No | Yes | Yes |
|  |  | TraesCS1B02G119300 | Yes | No | Yes | Yes |
|  |  |  | Yes | No | No | No |
| Starch synthase IIIb | *SSIIIb/SSIII-1* | TraesCS2A02G468800 | Yes | Yes | Yes | No |
|  |  | TraesCS2B02G491700 | Yes | No | Yes | No |
|  |  | TraesCS2D02G468900 | Yes | Yes | Yes | No |
| Starch synthase IV | *SSIVa/SSIV-1* | TraesCS1A02G353300 | Yes | No | Yes | No |
|  |  | TraesCS1B02G368500 | Yes | No | Yes | No |
|  |  | TraesCS1D02G356900 | Yes | Yes | Yes | No |

## Notes

*Declarations of interest*: none

https://www.ebi.ac.uk

